# Genomic selection for seed yield enhances flax breeding efficiency

**DOI:** 10.64898/2026.03.01.707406

**Authors:** Frank M. You, Chunfang Zheng, John Joseph Zagariah Daniel, Pingchuan Li, Kenneth Jackle, Megan House, Bunyamin Tar’an, Sylvie Cloutier

## Abstract

Genomic selection (GS) is a promising strategy to improve breeding efficiency for complex traits such as seed yield by enabling early selection and reducing reliance on extensive field testing. However, practical deployment of GS remains challenging due to limited training populations sizes and reduced predictive ability when models are applied to true breeding germplasm. In this study, we evaluated GS for flax (*Linum usitatissimum* L.) seed yield under realistic breeding scenarios, with a focus on across-population prediction (APP) and breeding decision support rather than model benchmarking. Using historical germplasm collections and a newly developed breeding-oriented population as training sets, GS performance was assessed across multiple independent test populations representing contemporary breeding lines evaluated in replicated yield trials. APP predictive abilities ranged from *r* = 0.67 to 0.84 depending on population relatedness when training and test populations were genetically aligned, supporting routine breeding deployment. Training population composition emerged as a key determinant of prediction success, with breeding-oriented populations consistently outperforming broad germplasm collections for predicting true breeding lines. Check-based selection analyses showed that GS reliably reproduced phenotypic advancement decisions while eliminating 61–91% of low-performing lines, resulting in 48–78% reduction in field evaluation costs for a typical cohort of 300 lines. Marker subsampling analyses further indicated that moderate-density genotyping-by-sequencing panels (∼2,500–3,000 SNPs) are sufficient to achieve stable predictive abilities. Overall, these results demonstrate that GS for seed yield in flax is ready for routine integration into breeding programs, offering a practical pathway to reduce costs, accelerate breeding cycles, and enhance selection efficiency.

## Introduction

Flax (*Linum usitatissimum* L.) is a dual-purpose crop cultivated for both oilseed (linseed) and fiber production, with distinct breeding objectives for each end use (Duk et al. 2021; Soto-Cerda et al. 2013; Povkhova et al. 2021; Praczyk and Wielgusz 2021). In linseed breeding programs, seed yield is the single most important selection target, followed by oil content and quality. As in all crops, genetic improvement for seed yield is challenging due to its complex genetic architecture, environmental sensitivity, and genotype-by-environment interactions. Consequently, phenotypic selection for YLD relies heavily on multi-year, multi-location field trials, making yield improvement both time-consuming and costly.

Genomic selection (GS), which uses genome-wide markers to predict genomic estimated breeding values (GEBVs), has emerged as a promising strategy to accelerate genetic gain for complex traits such as yield (Meuwissen et al. 2001; Olatoye et al. 2019; Beyene et al. 2019; Kumar et al. 2024). By enabling early-generation selection and reducing dependence on extensive field testing, GS has the potential to shorten breeding cycles and improve resource efficiency. In flax, GS has been explored for seed yield, agronomic traits, seed quality, and disease resistance (Hoque et al. 2024; Khan et al. 2023; Lan et al. 2020; He et al. 2018; You et al. 2022). Despite these advances, routine deployment of GS for seed yield in flax breeding programs remains limited, largely due to the small size of the training populations, shifting breeding pools, and uncertainties about the abilities of the prediction models to perform when applied to breeding germplasm unrelated to the training set(s).

A key limitation of many GS studies is their reliance on within-population cross-validation (CV) to assess predictive ability (PA). Because CV evaluates prediction within the same population used for model training, it often inflates PA. In practical breeding, however, GS models must predict untested breeding lines evaluated across different years and environments, a scenario more appropriately captured by across-population prediction (APP).Studies in cereals and legumes have shown that predictive abilities decline sharply under APP (Lozada et al. 2019; Juliana et al. 2018; Vitale et al. 2025), highlighting the need to evaluate GS strategies under conditions that reflect actual breeding deployments. Rather than benchmarking model architectures under idealized conditions, this study focuses on evaluating GS performance under deployment-oriented scenarios that mirror real breeding decision scenarios.

Beyond model choice, emerging evidence suggests that training population composition and genetic relevance to breeding targets are dominant factors governing APP performance. Training sets based on broad germplasm collections provide extensive diversity but often include genetic variation that is weakly informative for predicting elite breeding germplasm. In contrast, breeding-oriented training sets composed of recent selections and parental lines better represent the genetic architecture of future candidates, thereby improving prediction robustness. However, systematic evaluations of training set designs, marker representations, and practical decision outcomes for seed yield prediction in flax are still lacking.

The objectives of this study were therefore to evaluate GS for flax seed yield under actual breeding scenarios, with an emphasis on APP and breeding decision support rather than model benchmarking. Specifically, we aimed to (i) assess how training set design influences PA across independent breeding populations, (ii) compare GS performance of multiple marker representations under APP conditions, (iii) determine the minimum marker density required for reliable prediction of breeding lines, and (iv) evaluate the practical utility of GS for check-based selection decisions, and in terms of cost reduction and breeding cycle acceleration.

By integrating historical germplasm collections with newly developed breeding populations and evaluating GS performance across multiple independent test datasets, this study provides a deployment-oriented assessment of GS for seed yield in flax. Our results offer practical guidance for training population construction, genotyping strategy, and GS integration into routine flax breeding pipelines.

## Materials and methods

### Training populations

A core collection was originally assembled to capture the genetic diversity of the global flax germplasm (Diederichsen et al. 2013). The initial collection comprised 381 accessions from 38 countries, including fiber and linseed landraces, breeding lines, and cultivars (You et al. 2017). To better represent modern breeding germplasm, 10 breeding lines and cultivars from Canadian breeding programs were incorporated, expanding the core collection to 391 accessions. The final set included 298 linseed and 93 fiber accessions, of which 378 lines (293 linseed and 85 fiber) had complete phenotypic datasets. Two GS training sets based on the core collection were defined: CORE293, consisting of 293 linseed accessions, and CORE378, comprising 378 accessions representing both linseed and fiber flax.

To better represent contemporary breeding germplasm and increase genetic diversity and relatedness to current breeding materials, a breeding-oriented training population, designated BP295, was developed. BP295 contains two germplasm components. The first component included 45 varieties and breeding lines that have been used as parents in recent years in the Crop Development Centre (CDC, University of Saskatchewan, Canada) flax breeding program. This set included cultivars released within the past 15 years, which were largely absent from the original core collection, as well as germplasm recently introduced from foreign breeding programs. Because these accessions have been extensively used as parental lines, they provide strong genetic connectivity to contemporary breeding populations. These 45 parental lines were not phenotyped in the same historical trials as the CORE populations and therefore could not be directly merged with the core collection. To generate a unified dataset, 254 breeding lines derived from recent breeding populations were selected in 2023. These 254 lines and the 45 parental accessions were evaluated together in the same field experiments conducted in 2025, ensuring directly comparable phenotypic data. After quality control, 295 of 299 lines with complete genotypic and phenotypic data were retained, forming the BP295 dataset, which was used as both a training and a test dataset in GS analyses.

### Test populations

In addition to BP295, three independent test populations were used for APP. The first test population comprised a combined set of 260 lines (255 inbred lines plus five of the six parents (excluding the reference cultivar CDC Bethune) from three biparental populations (BM, EV, and SU) with distinct genetic backgrounds (You et al. 2018a). The BM population comprised 91 F_6_-derived recombinant inbred lines (RILs) from a cross between the high-yielding cultivars CDC Bethune and Macbeth. The EV population consisted of 88 F_6_-derived RILs from a cross between the low linolenic acid breeding line E1747 and the French fiber cultivar Viking. The SU population contained 76 F_1_-derived doubled haploid (DH) lines from a cross between the yellow-seeded, low linolenic acid breeding line SP2047 and the high linolenic acid breeding line UGG5-5.

The second and third test populations consisted of yellow-seeded (YS) and brown-seeded (BS) breeding lines, targeting the newer yellow-seeded and the traditional, brown-seeded market classes, respectively. These lines were selected from the CDC’s flax breeding program in 2023 and 2024. A total of 33 YS lines and 57 BS lines, together with their respective parent and check cultivars, were evaluated in four independent yield trials conducted across three or four locations during 2023–2024 using randomized complete block designs (RCBD) with three replications per location (Table 1). For GS analyses, the YS lines, associated parents and check cultivars were designated YS38, and the similar BS set was designated BS61.

**Table 1.** Summary of yield trials used for genomic prediction of seed yield.

| Yield trial | Year | Code<sup>†</sup> | No. of test lines | Location | Check/parent cultivars<sup>‡</sup> |
| --- | --- | --- | --- | --- | --- |
| Yellow seed secondary | 2024 | 2024YSS (YS38) | 28 | U of S Campus, Kernen, Melfort | **AAC Bright (2017)**, CDC Dorado (2016), CDC Melyn (2018), and Double-Low |
| Yellow seed secondary and tertiary | 2023–2024 | 2023–24YST (YS38) | 5 | U of S Campus, Kernen, Melfort, Avonlea (2024 only) | **AAC Bright (2017)**, CDC Dorado (2015), and NuLin VT50 (2009) |
| Brown seed secondary | 2024 | 2024BSS (BS61) | 37 | U of S Campus, Kernen, Melfort | CDC Kernen (2021), CDC Esme (2023), **CDC Glas (2012)** |
| Brown seed secondary and tertiary | 2023–2024 | 2023–24BST (BS61) | 20 | U of S Campus, Kernen, Melfort, Avonlea (2024 only) | CDC Kernen (2021), CDC Esme (2023), **CDC Glas (2012)**, and CDC Rowland (2018) |
| Preliminary | 2025 | BP295 | 299 | U of S Campus | **CDC Glas (2012)**, **AAC Bright (2017)**, CDC Bethune (1998), CDC |
|  |  |  |  |  | Kernen (2021), CDC Plava (2015) |

<sup>†</sup> Codes indicate the experiment, with test population codes shown in parentheses. <sup>‡</sup> National check cultivars (in bold) are AAC Bright for yellow-seeded, and CDC Glass for brown-seeded trials. CDC Rowland is an internal check. Numbers in parentheses denote years of cultivar release. U of S: University of Saskatchewan

### Sequencing and single nucleotide polymorphism (SNP) markers

All individuals of the flax core collection (CORE378), the combined biparental population (BMEVSU260) and the 45 varieties and breeding lines of BP295 were genotyped by whole-genome sequencing (WGS) on Illumina platforms using the 100- (PE100) or 150-bp (PE150) paired end read mode, as previously described (You et al. 2018a; He et al. 2023). The 99 breeding lines and check/parent cultivars from the CDC flax breeding program (YS38 and BS61) and the 254 breeding lines of the BP295 set were genotyped using genotyping-by-sequencing (GBS) with PE150 reads using the *Pst*I–*Msp*I enzyme protocol (Elshire et al. 2011).

All lines were grown in root trainers (72 plants per tray) in a growth chamber under a 20 h light/4 h dark cycle (24°C/18°C). When seedlings were 5-8 cm tall, approximately 50–80 mg of leaf tissue was collected from the shoot tip into Qiagen 96-well collection tubes pre-cooled on dry ice. Samples were lyophilized for 48-72 h in a FreeZone 6 L benchtop freeze dryer (Labconco, Kansas City, MO, USA) and stored at −20°C until DNA extraction.

For GBS, genomic DNA (gDNA) was extracted with the Qiagen DNeasy® kit (Qiagen, Toronto, ON, Canada) and quantified using the Quant-iT™ PicoGreen™ dsDNA assay (ThermoFisher Scientific, Waltham, MA, USA) following the manufacturers’ instructions. The gDNA was normalized to 10 ng/µl, and 200 ng samples were submitted to the Université Laval’s Institut de biologie intégrative et des systèmes (IBIS; Quebec, QC, Canada) for library preparation. GBS libraries were pooled and sequenced on the Illumina HiSeq 2500 platform to generate PE150 reads, as previously described in Cloutier et al. (2024).

For SNP discovery, all reads were aligned to the CDC Bethune v3.0 reference genome (You et al. 2026a) using bwa v0.7.19 (Li and Durbin 2009). Single nucleotide polymorphisms (SNPs) were called with SAMtools v1.22.1 (Li et al. 2009) and filtered using the following threshol criteria: read depth > 3, mapping quality (MAPQ) on the Phred scale > 20, minor allele frequency (MAF) ≥ 0.05, call rate ≥ 80%, heterozygosity ≤ 20% (per SNP), and exclusion of multi-allelic sites. The workflow was implemented in the improved AGSNP pipeline (Kumar et al. 2012; You et al. 2012). After quality control using the above criteria, SNPs in all training and test datasets were imputed using Beagle v5.5 with default parameters (Browning et al. 2021). The distribution of filtered SNPs was visualized using the CMplot R package (v4.5.1) with a 100-Kb window size (Yin et al. 2021).

### Haplotype and PC marker construction

In addition to SNP, haplotype block (HAP) and principal component (PC) markers were generated. HAPs were estimated with the rtm-gwas-snpldb tool in RTM-GWAS v2020.0 (He et al. 2017), which applies the Haploview “Gabriel” confidence-interval algorithm (Gabriel et al. 2002; Barrett et al. 2005). Pairwise linkage disequilibrium (LD) was measured using D′ with 95% confidence intervals. SNP pairs were classified as strong LD, inconclusive, or strong recombination, and HAPs were defined when ≥95% of informative pairs were in strong LD (default thresholds: CI(D′) lower ≥ 0.70, upper ≥ 0.98). PC markers were obtained by principal component analysis (PCA) of the SNP dataset when the first *N* PCs explaining 95% of total variance were retained for prediction.

### Yield data analysis

The core collection (378 accessions) was phenotyped for seed yield over four years (2009–2012) at two locations: the Morden Research and Development Centre Farm, Morden, Manitoba, and the Kernen Crop Research Farm, near Saskatoon, Saskatchewan, Canada (You et al. 2017). The experimental design was a type-2 modified augmented design (MAD2) (Lin and Poushinsky 1985). Yields were adjusted for soil heterogeneity using the MAD pipeline (You et al. 2013). Best linear unbiased predictions (BLUPs) across years and locations were obtained using *lme4* v1.1-37 (Bates et al. 2015), with genotypes, years and locations treated as random effects.

The BM, EV and SU populations were phenotyped at the same locations. BM was evaluated for four years (2009–2012), and EV and SU for three years (2010–2012), also using MAD2 designs. Phenotypic adjustments and BLUP estimation followed the same procedure as the core collection (You et al. 2013).

The 90 selected breeding lines and nine checks/parents making out the test populations were evaluated in independent yield trials in 2023–2024 at three (secondary trials) or four (tertiary trials) locations (Table 1). Linear mixed models were fitted using test lines (genotypes) as fixed effects, and blocks (replications), trials, locations, years, and lines × locations as random effects. The yield estimates of lines measured through the mixed model were considered observed (“true”) breeding values.

Field evaluation of the BP295 population was conducted using a structured multi-experiment design to control field heterogeneity and improve phenotypic reliability. The 296 lines were divided into eight subsets, each containing 37 lines. Each subset was evaluated as an independent experiment using RCBD with two replications. Five common check cultivars were included in every experiment, resulting in 42 entries per experiment and a total of eight parallel experiments. To further assess data robustness and minimize potential seed source effects, a second, independent set of eight experiments was conducted in a separate field using the same experimental design and line allocation. The only difference between the two experimental sets was the seed source: one set used seeds increased in a greenhouse at AAFC (Ottawa, Canada), and the other used seeds provided by the CDC which were advanced in New Zealand during the winter of 2024. For both seed sources, seed increases were derived from a single plant per line to ensure genetic uniformity.

Phenotypic data from the two experimental sets were first analyzed separately using a two-stage multi-environment trial (MET) approach (Piepho et al. 2012; Piepho et al. 2008). In the first stage, adjusted least squared means were obtained for each line within individual experiments using mixed linear models accounting for block effects. In the second stage, line effects were jointly analyzed across experiments to obtain overall seed yield estimates. Seed yield estimates derived from the two experimental sets were highly correlated (Pearson’s *r* = 0.85), indicating strong consistency between seed sources. Consequently, data from both sets were combined, and a final two-stage MET analysis was performed to generate consolidated BLUPs for seed yield, which were used in subsequent GS analyses.

### Genomic selection models and predictive ability evaluation

Sixteen GS models implemented in the MultiGS-P pipeline (You et al. 2026b) were evaluated, representing a range of statistical, machine learning (ML), and deep learning (DL) approaches. These included four linear and regularized models (R_RRBLUP, R_GBLUP, Elastic Net, and BRR), three ML models (Random Forest Regression (RFR), XGBoost and LightGBM), six DL models, two hybrid model that integrate RR-BLUP with deep networks (DeepResBLUP and DeepBLUP), and a stacking-based ensemble learner (EnsembleGS).

The DL models comprised multilayer perceptron-based architectures (MLPGS and DNNGS) and graph-based neural networks (GraphConvGS, GraphAttnGS, GraphSAGEGS, and GraphFormer), designed to capture nonlinear genotype–phenotype relationships and exploit marker dependencies. EnsembleGS combined predictions from multiple base models (R_RRBLUP, Elastic Net, BRR, LightGBM, and MLPGS) using a stacking strategy to enhance robustness and predictive performance. These models were selected to represent GS approaches of increasing complexity and flexibility.

Default hyperparameters, which were carefully tuned using the flax datasets and provided by MultiGS-P (You et al. 2026b), were applied to all ML and DL models. Key hyperparameters for deep learning models are summarized in Table S1. For each model, SNP, HAP, and PC marker representations were evaluated.

Predictive ability (PA) was defined as the Pearson correlation between genomic estimated breeding values (GEBVs) and observed phenotypic values. Because phenotypic values include environmental variance, PA reflects predictive ability rather than the true accuracy of breeding values. Two evaluation strategies were employed: (i) five-fold CV within training populations, repeated across 10 training-test partitions (yielding 50 total CV replicates, as each partition generated five folds used in turn as the test set), to assess model and marker-representation performances, and (ii) APP, in which three training populations were used to predict four independent breeding line datasets.

### Evaluation of predictive abilities across marker numbers in across-population prediction

To determine the number of SNP markers required for reliable prediction of new breeding lines, simulations were conducted using randomly sampled SNP subsets. For the BMEVSU260 test dataset, 33,596 SNPs shared between the test and training populations were available, from which subsets ranging from 500 to 10,000 SNPs (in increments of 500) were generated, with ten random replications per subset. The rationale for capping the SNPs at 10,000 was two-fold: (i) preliminary analyses had demonstrated that PA reached a plateau when marker density exceeded that number, and (ii) the aim was to determine the minimum number of markers required for practical APP. For the BP295 test population, approximately 6,000 SNPs were common between the test and training populations, and subsets ranging from 500 to 5,500 SNPs (in increments of 500) were similarly generated, with ten replications per subset.

The representative GS models R_RRBLUP, LightGBM, and DeepBLUP, implemented in MultiGS-P, were evaluated using SNP markers. PAs were then compared across incremental marker numbers and models to assess the trade-off between marker density and APP performance.

### Population genetic diversity and differentiation analysis

To assess genetic diversity and relatedness of training and test populations, nucleotide diversity within populations and genetic differentiation between populations were quantified using harmonized SNP markers. For each population, SNP-based nucleotide diversity (π) was calculated as the mean pairwise nucleotide difference per SNP. Prior to analysis, SNPs were harmonized across all VCF files of training and test datasets to retain only shared loci with consistent reference and alternate alleles.

Pairwise population genetic differentiation was estimated using the Weir–Cockerham *F_ST_* statistic (Weir and Cockerham 1984) based on shared SNPs between population pairs. Genotype data from each pair of populations were combined, and subpopulation assignments were used to calculate *F_ST_* components following the original Weir–Cockerham formula. All diversity and differentiation statistics were computed using a custom Python pipeline built on the scikit-allel framework, which is implemented in the MultiGS toolkit (https://github.com/AAFC-ORDC-Crop-Bioinfomatics/MultiGS).

### Economic and cost–benefit analysis

A cost–benefit analysis was performed to compare conventional phenotypic selection with GS under practical flax breeding scenarios. Cost parameters were derived from routine operations in the CDC flax breeding program and included field evaluation, laboratory phenotyping, and genotyping expenses. Field costs were estimated on a per-plot basis and scaled to per-line costs according to the number of replications used in preliminary yield trials. Additional per-line costs for routine phenotyping (e.g., oil profile analysis and disease screening) were incorporated.

Genotyping costs were estimated for GBS, including DNA extraction, library preparation, and sequencing. Sequencing costs were modeled as a function of read depth and sample multiplexing based on NovaSeq platform specifications we used. Representative sequencing depth and marker density were defined based on marker subsampling analyses (see Results), which informed the minimum number of SNPs required for stable predictive ability.

Total breeding costs under GS-assisted selection were calculated by combining genotyping costs with reduced field evaluation costs, where the number of lines advanced to field trials was adjusted according to genomic prediction-based pre-selection. Cost savings were estimated by comparing total expenditures between GS-assisted and conventional phenotypic selection scenarios.

## Results

### Training and test datasets

A total of six flax populations were included in this study, comprising three training populations and four test populations that collectively represent both historical germplasm collections and contemporary breeding germplasm (Table 2).

**Table 2.** Summary of training and test populations used in flax genomic selection.

| Population type | Population description | Code | Lines with trait data | Total filtered SNPs <sup>†</sup> | Common SNPs used for prediction<sup>‡</sup> | Nucleotide diversity ($\pi$) |
| --- | --- | --- | --- | --- | --- | --- |
| Training | Core collection | CORE378 | 378 | 1,699,363 | 5,604 | 0.3991 |
|  | Linseed subset of the core collection | CORE293 | 293 | 1,696,929 | 5,604 | 0.4007 |
| Training/Test | Breeding population (252 breeding lines + 43 parents) | BP295 | 252 | 11,235 | 5,604 | 0.3838 |
|  |  |  | 43 | 1,218,703 | 5,604 |  |
| Test | Combined biparental population | BMEVSU260 | 260 | 43,268 | 33,596 | 0.3727 |
|  | Yellow-seeded breeding lines | YS38 | 38 | 11,714 | 5,604 | 0.3549 |
|  | Brown-seeded breeding lines | BS61 | 61 | 11,714 | 5,604 | 0.2891 |

<sup>†</sup> Number of SNPs after filtering using the following criteria: SNP call rate $\geq 0.8$ , heterozygous rate $\leq 0.2$ , and minor allele frequency $\geq 0.05$ . YS and BS were analyzed together as a single group for SNP discovery but evaluated separately for predictive ability. <sup>‡</sup> The value 33,596 denotes SNPs shared between BMEVSU260 and the CORE378 and CORE293 datasets, where 5,604 denotes SNPs shared across all training datasets and the YS38 and BS61 test datasets

All accessions in the CORE populations (CORE378 and CORE293) were genotyped using whole-genome shotgun sequencing (WGS) (You et al. 2018a; He et al. 2023). Following SNP discovery and filtering, 1,699,363 high-quality SNPs were retained in CORE378 and 1,696,929 SNPs in CORE293 (Fig. S1A). These populations exhibited high nucleotide diversity (π = 0.3991–0.4007), consistent with their broad and diverse germplasm origins.

In BP295, the 252 breeding lines were genotyped using a GBS platform, generating an average sequence depth of 2.5 Gb per line and a mean coverage of 5.16× (Table S2). The 43 parental or germplasm accessions were genotyped using WGS, and SNPs common to both datasets were retained. After harmonization, 5,604 SNPs were shared across all BP295 lines (Fig. S1C). Despite its breeding-focused composition, BP295 displayed nucleotide diversity (π = 0.3838) comparable to that observed in the CORE populations. Owing to its genetic relevance to contemporary breeding germplasm, BP295 was evaluated both as an independent training population and, in some analyses, as a test population. When BP295 was used as a test population in APP analyses, the training data consisted exclusively of the CORE293 or CORE378 populations, no data from BP295 was used in model training for these specific evaluations.

The lines from the test population BMEVSU260 were genotyped using WGS, generating an average of more than 28 million PE100 reads per sample and a mean genome coverage of 11.41× (Table S2). Of these, 94.47% of reads aligned to the CDC Bethune v3.0 reference genome, yielding 43,268 high-quality SNPs (You et al. 2018b), of which 33,596 overlapped with the CORE training populations (Fig. S1B; Table 2).

For the second and third test populations (YS38 and BS61) (Table 1), GBS produced an average of 6.0 million PE150 reads per sample, with 97.70% of reads mapping to the CDC Bethune v3.0 reference genome (Table S2). This resulted in 11,714 high-quality SNPs, of which 5,604 were shared with the training populations (Fig. S1C; Table 2).

Due to differences in genotyping platforms, the number of SNPs shared between training and test populations varied. BMEVSU260 shared 33,596 SNPs with the CORE populations but only ∼300 SNPs with GBS-based datasets, whereas YS38, BS61, and BP295 shared 5,604 SNPs, making them more directly comparable for APP analyses.

Pairwise population genetic differentiation, estimated using Weir–Cockerham *F_ST_* (Weir and Cockerham 1984) (Table 3), revealed clear contrasts among populations. The CORE populations showed moderate to strong differentiation from the breeding test populations (*F_ST_* = 0.2458–0.3739), reflecting their broader and more historical germplasm composition. In contrast, BP295 exhibited lower differentiation from YS38 (*F_ST_* = 0.0899) and BS61 (*F_ST_* = 0.1323) than either CORE293 or CORE378, indicating closer genetic proximity to contemporary breeding germplasm. BMEVSU260 showed moderate differentiation from the CORE populations (*F_ST_* ∼0.24-0.26), consistent with its biparental origin. Collectively, these results demonstrate that BP295 occupies an intermediate genetic position between historical germplasm collections and modern breeding populations, supporting its use as a biologically relevant training population for genomic prediction in applied flax breeding.

**Table 3.** Pairwise population genetic differentiation (*F_ST_*) among training and test populations.

| Population | CORE293 | CORE378 | YS38 | BS61 |
| --- | --- | --- | --- | --- |
| CORE378 | 0.2458 |  |  |  |
| YS38 | 0.2574 | 0.2712 |  |  |
| BS61 | 0.3488 | 0.3739 | 0.2036 |  |
| BP295 | 0.2379 | 0.2548 | 0.0899 | 0.1323 |

### Cross-validation based predictive ability for seed yield

CV was first used to evaluate the prediction capacity of different training populations, genomic selection models, and marker representations under ideal within-population conditions. PA, measured as the Pearson correlation between predicted and observed YLD, was assessed using a five-fold CV scheme with ten replicates (50 runs in total) for each combination of training population, model, and marker representation (Fig. 1; Table S3).

**Fig. 1.**
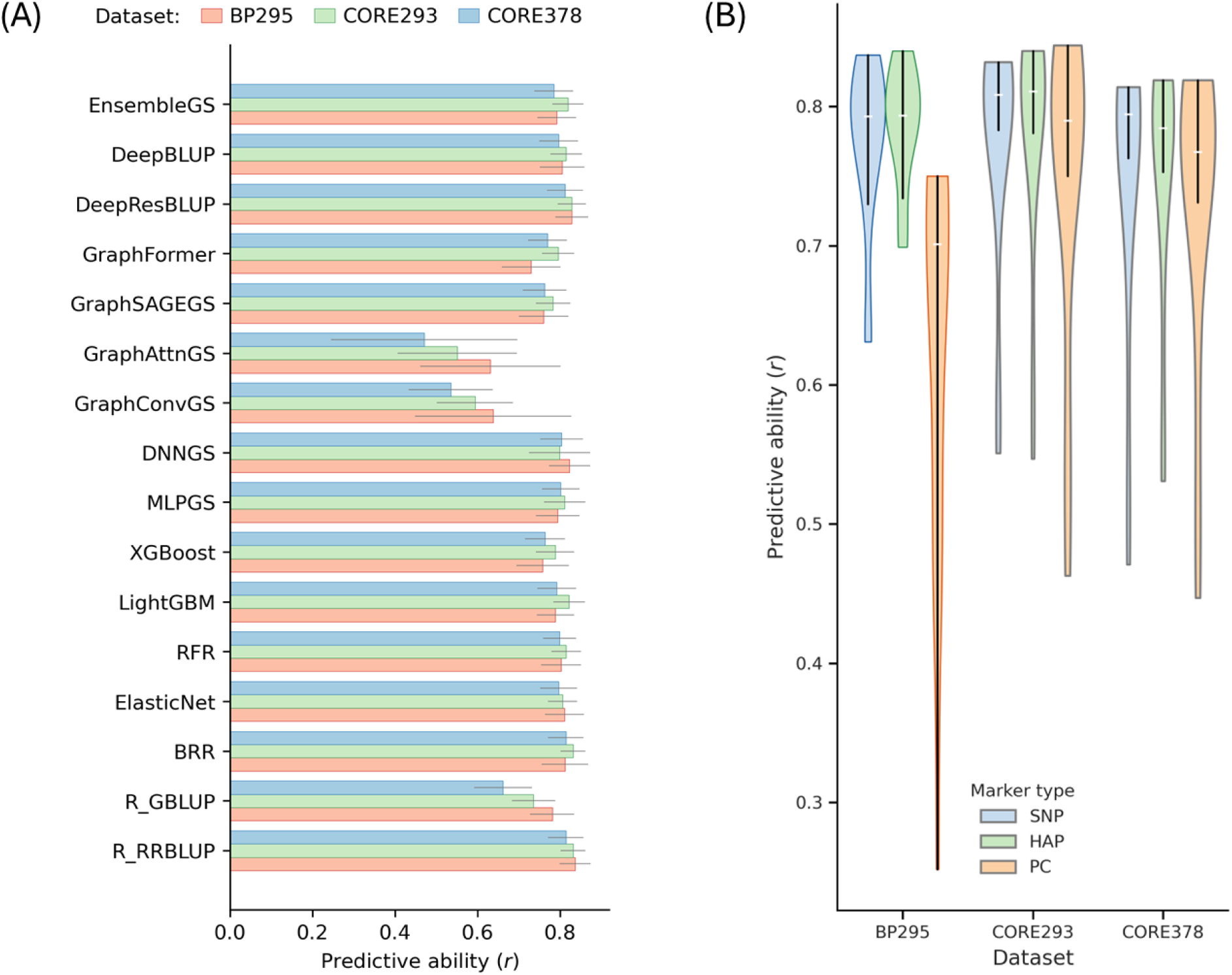
Cross-validation (CV)–based predictive abilities (PAs) for seed yield. (A) Bar chart showing the mean PA for each model using SNP markers, with error bars indicating the standard deviation. (B) Violin plots summarizing the distribution of PAs across 16 models and all three marker representations. PA was evaluated using a five-fold CV scheme with 10 random training-test partitions, resulting in 50 total replicates for each training dataset

Across the three training populations, CV-based PAs were generally high, with mean values typically ranging from 0.75 to 0.85 for most linear, ML, and DL models when using SNP or HAP markers (Table S3). The highest mean PA observed across all CV analyses was 0.84, achieved by RR-BLUP and BRR models using BP295 or CORE293 as the training population. In contrast, predictive abilities based on PC representations were consistently lower and exhibited higher variability across models, particularly for DL architectures (Fig. 1B; Table S3).

When comparing training populations, BP295 showed CV performance comparable to or slightly exceeding that of CORE293 and CORE378 across most models and marker representations. Notably, BP295 maintained high PAs despite being composed primarily of recent breeding lines rather than broad germplasm accessions, indicating that its genetic diversity is sufficient to support effective within-population prediction. CORE293 generally outperformed CORE378 across models, whereas CORE378 exhibited a modest reduction in PA for several models, consistent with its mixed linseed and fiber flax composition.

Model-specific trends were largely consistent across training populations. Linear mixed models (RR-BLUP, GBLUP and BRR) performed robustly, achieving high and stable PAs with both SNP and HAP markers. Tree-based ML models (RFR, XGBoost and LightGBM) and shallow DL models (MLPGS and DNNGS) performed comparably, with no systematic advantage over linear baselines under CV conditions. Among graph-based DL models, GraphSAGEGS and GraphFormer consistently outperformed GraphConvGS and GraphAttnGS, although their overall performance remained slightly below that of the best linear and hybrid models. Hybrid approaches integrating RR-BLUP with deep architectures (DeepBLUP and DeepResBLUP) consistently achieved predictive abilities comparable to, and in some cases marginally higher than, linear mixed models, while exhibiting moderate variability across training populations.

The distribution of CV-based PAs across all 16 models further highlighted differences among marker representations (Fig. 1B). SNP and HAP markers produced similarly high and stable PA distributions across training populations, whereas PC-based markers resulted in broader and lower PA distributions, particularly for BP295. These results indicate that dimensionality-reduced PC features are less effective at capturing the genetic signal underlying seed yield under CV conditions, especially when training populations consist of heterogeneous breeding germplasm.

Overall, CV-based analyses demonstrated that multiple model classes and marker representations can achieve high PA under ideal within-population conditions, and that BP295 performs at least as well as historical CORE collections as a training population. However, because CV relies on random partitions within the same population, the results primarily reflect model capacity rather than predictive robustness in real-life breeding scenarios. Consequently, APP analyses were conducted to evaluate model performance under conditions that emulate practical breeding deployment.

### Across-population predictive ability for seed yield

Four independent test datasets—BMEVSU260, YS38, BS61, and BP295—were predicted using three marker representations (SNP, HAP and PC), with models trained on the historical core collections (CORE378 and its linseed-only subset CORE293) and, where applicable, the breeding-oriented training population BP295.

Across all test datasets, APP-based PAs were generally lower and more variable than those observed under cross-validation (Fig. 2, Tables S4-S6), reflecting the increased difficulty of predicting across genetically and temporally distinct populations. Nevertheless, PAs frequently exceeded 0.60 and, in some cases, reached 0.80 or higher, demonstrating the feasibility of genomic prediction for seed yield under actual breeding schemes.

**Fig. 2.**
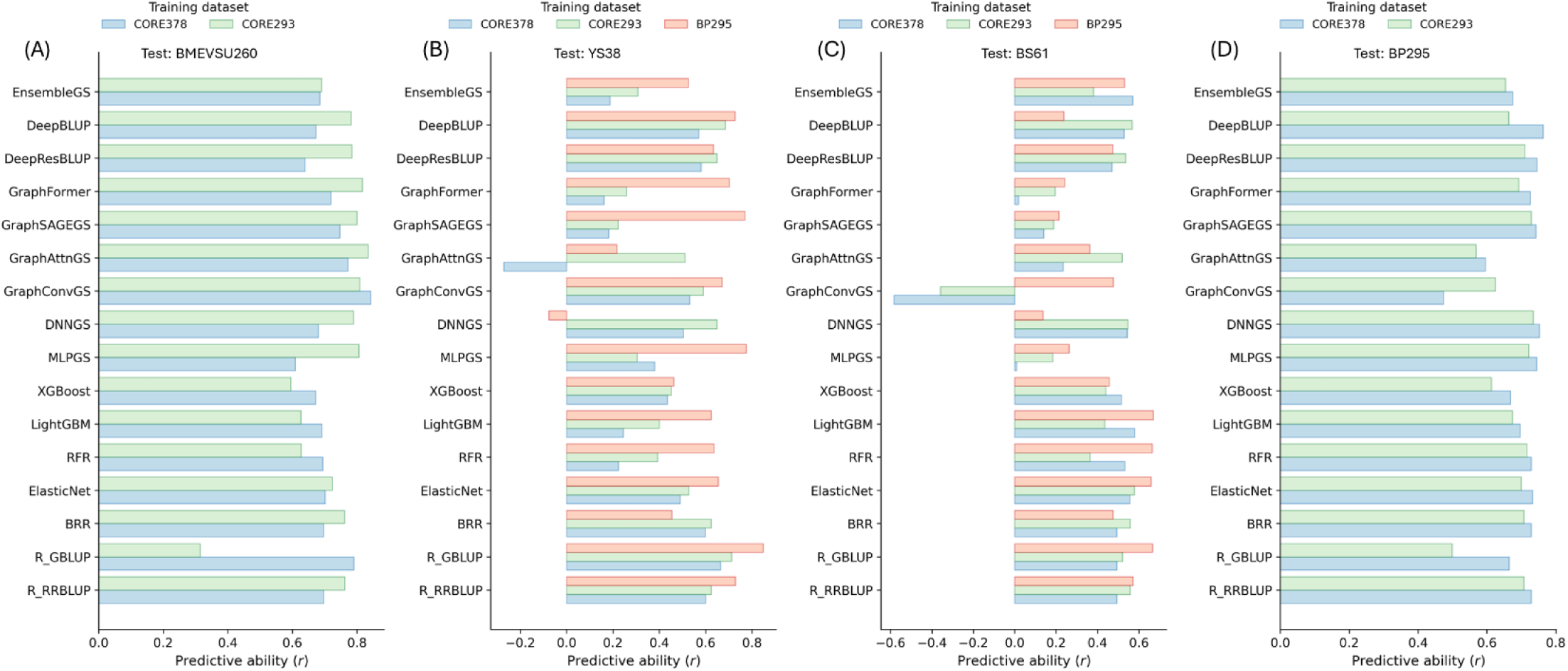
Predictive abilities (PAs) of seed yield obtained using 15 genomic selection models with SNP markers across different training populations and test datasets: (A) BMEVSU260, (B) YS38, (C) BS61, and (D) BP295

Training population choice had a pronounced impact on APP performance. In particular, BP295 consistently outperformed the historical CORE collections when used as the training population for predicting contemporary breeding lines. For the YS38 test set, BP295 achieved the highest PA for most models, with maximum values reaching 0.85 with GBLUP, exceeding the PA values obtained using CORE293 or CORE378 (Fig. 2B). Similar trends were observed for BS61, where BP295 generally provided higher or comparable PAs relative to CORE-based training, despite the greater variability observed across models for this training set (Fig. 2C).

In contrast, for the BMEVSU260 biparental population, CORE293 consistently outperformed CORE378 across nearly all models, with maximum PAs reaching 0.84 for several graph-based and hybrid models (Figs. 2A and 3A). This result is consistent with the close genetic relationship between BMEVSU260 and the linseed-only CORE293 population (Table 2). BP295 was not evaluated as a training population for BMEVSU260 due to limited marker overlap resulting from differences in genotyping platforms.

**Fig. 3.**
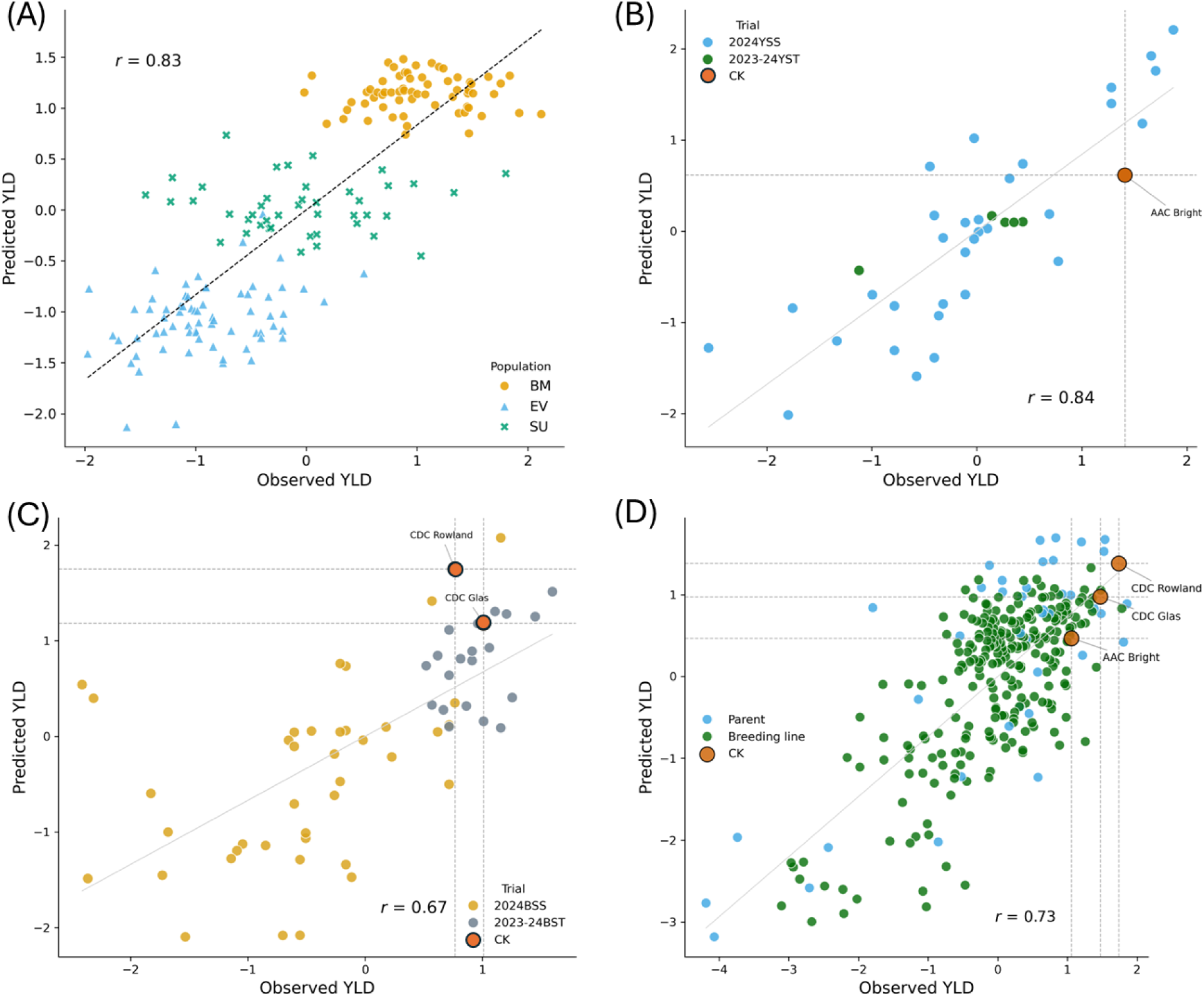
Across-population prediction (APP) of seed yield (YLD) using four test datasets illustrating the ability of genomic selection to reproduce check-based advancement decisions while filtering effectively low-performing lines. (A) Scatter plot of observed versus predicted seed yield for 260 combined biparental inbred lines (BMEVSU260) using the MLPGS model with 7,871 haplotype (HAP) markers. (B) Scatter plot of observed versus predicted seed yield for the YS38 dataset using the GBLUP model with 1,945 HAP markers. (C) Scatter plot of observed versus predicted seed yield for the BS61 dataset using the LightGBM model with 1,933 HAP markers. (D) Scatter plot of observed versus predicted seed yield for the BP295 dataset using the deep learning model DNNGS with 5,604 SNP markers. “CK” denotes check cultivars. Vertical and horizontal reference lines indicate thresholds for check-based selection under genomic and phenotypic selection. Observed and predicted see yield values were standardized to a mean of 0 and a standard deviation of 1

When BP295 was treated as test population, CORE378 achieved slightly higher predictive abilities (0.01–0.03) than CORE293 across models (Fig. 2D), despite marginally higher genetic differentiation between CORE378 and BP295 (*F_ST_* = 0.2548) compared with CORE293 and BP295 (*F_ST_* = 0.2349). This pattern likely reflects the broader allelic diversity and genetic coverage of CORE378, which includes both linseed and fiber germplasm and therefore more fully represents the genetic space occupied by BP295 (Fig. S2). In contrast, CORE293 contains only linseed accessions and captures a narrower allelic spectrum. These results indicate that, when the test population lies largely within the diversity range of the training population, broader genetic coverage can outweigh slight differences in genetic relatedness in determining PA using APP.

Across all APP analyses, no single model class consistently dominated all test datasets, and relative model rankings differed from those observed under CV. Linear mixed models (RR-BLUP, GBLUP, BRR) and regularized regression models (ElasticNet) showed robust and stable performances across most scenarios, frequently ranking among the top-performing methods. Tree-based ML models (Random Forest, XGBoost, LightGBM) exhibited moderate to high PA, although their performance was more sensitive to training population choice.

DL and hybrid models displayed a wide range of performances under APP. Hybrid models integrating RR-BLUP with deep architectures (DeepBLUP and DeepResBLUP) consistently achieved high PAs across multiple test datasets, particularly when training and test populations were genetically compatible. Graph-based DL models showed robust performance for BMEVSU260, with GraphConvGS and GraphAttnGS achieving some of the highest PAs (> 0.83), but their performance was less stable for YS38 and BS61, occasionally yielding low or even negative correlations when training–test divergence was large. These results indicate that model complexity alone does not guarantee superior APP performance, and that robustness across populations remains a critical consideration.

Marker representation influenced APP performance, although its effect was generally secondary to training population choice. Across test datasets, SNP- and HAP-based markers consistently outperformed PC-based markers, particularly for BP295-trained models predicting YS38 and BS61. PCs frequently resulted in reduced or unstable PAs, especially for graph-based DL models. Nevertheless, the relative performance of SNPs and HAPs was broadly comparable across models, suggesting that moderate-density GBS markers are sufficient for effective APP of seed yield when appropriate training populations are used.

Taken together, APP analyses demonstrate that training population composition is a primary determinant of PA for seed yield in flax, often outweighing differences among models or marker representations. The breeding-oriented population BP295 consistently provided superior or comparable prediction performance for contemporary breeding lines, whereas historical CORE collections performed best when predicting genetically related materials. These results highlight the importance of aligning training population design with breeding objectives and confirm the merit of APP-based evaluation for evaluating GS strategies intended for practical deployment.

### Check-based selection decision of breeding lines

Because breeding decisions are ultimately threshold-based rather than correlation-based, we further evaluated GS performance using check-based advancement decisions. To evaluate the practical utility of GS for breeding decision-making, we examined GS-based predictions for the YS38, BS61 and BP295 test datasets using check cultivars as benchmarks for advancement decisions. In Canadian flax breeding programs, AAC Bright is the main check cultivar for yellow-seeded flax, whereas CDC Glas and CDC Rowland (advanced trials only) serve as standard checks for brown-seeded flax. For each test dataset, the training population and model combination that achieved the highest APP predictive ability was used for CK-based selection analyses.

For the YS38 dataset, predictions generated using BP295 as the training population and the GBLUP model with HAPs achieved the highest PA (*r* = 0.84). Using AAC Bright as the benchmark check, GS accurately identified all four breeding lines that outperformed the check cultivar for observed seed yield, together with four additional promising candidates that were not selected based solely on phenotypic performance (Fig. 3B). Importantly, GS consistently excluded five lines from the 2023–2024 YS trial that underperformed the check cultivar in both observed and predicted yields (green dots in Fig. 3B). If GS-based selection had been used, the number of lines advanced to subsequent trials would have been reduced from 33 to 8, representing a 75% reduction in field-testing requirements, while retaining all superior-yielding candidates identified by phenotypic selection.

For the BS61 dataset, the highest predictive ability was achieved using BP295 as the training population with the LightGBM model and haplotype markers (*r* = 0.67). Using CDC Glas as the primary benchmark check (Fig. 3C), GS identified four of the seven lines exceeding the check based on observed yields (57%), along with one additional high-GEBV candidate. The three GS-missed superior lines ranked third, fifth, and seventh phenotypically, indicating low risk of excluding elite candidate lines. Under this GS-assisted decision framework, only five lines would be advanced, representing a 91% reduction in trial size (from 57 to 5).

For the BP295 dataset, analyses focused on the 252 new breeding lines, excluding parental and germplasm accessions. Using CORE293 as the training population, the DNNGS model with SNP markers achieved the highest PA (*r* = 0.73). When AAC Bright was used as the benchmark check, phenotypic selection identified 15 lines superior to the check based on predicted breeding values. Applying GS-based selection, 95/252 lines would be advanced to the next stage of yield testing, representing a 62% reduction. Among these, 13/15 lines (87%) selected based on their phenotypic performance would be retained, including the two highest-yielding lines across all replicated field trials. GS-based selection can reduce population size for advanced testing while maintaining a minimal risk of excluding elite candidate lines.

Across all three test datasets, GS effectively reproduced check-based phenotypic advancement decisions while consistently filtering out low-performing entries. By enabling earlier selection, reducing the number of lines prior to extensive multi-environment testing, and preserving high-performing candidates, GS has the potential to enhance breeding efficiency and accelerate cultivar development in flax.

### Number of markers required for prediction of breeding lines

To assess the minimum marker density required for effective genomic prediction of breeding lines, across-population prediction analyses were conducted using random subsets of SNPs for BMEVSU260 and BP295, with CORE293 as the training population. Three models representing linear, machine-learning, and hybrid approaches (R_RRBLUP, LightGBM, and DeepBLUP) were evaluated using increasing numbers of randomly sampled SNPs shared between training and test populations (Fig. 4).

**Fig. 4.**
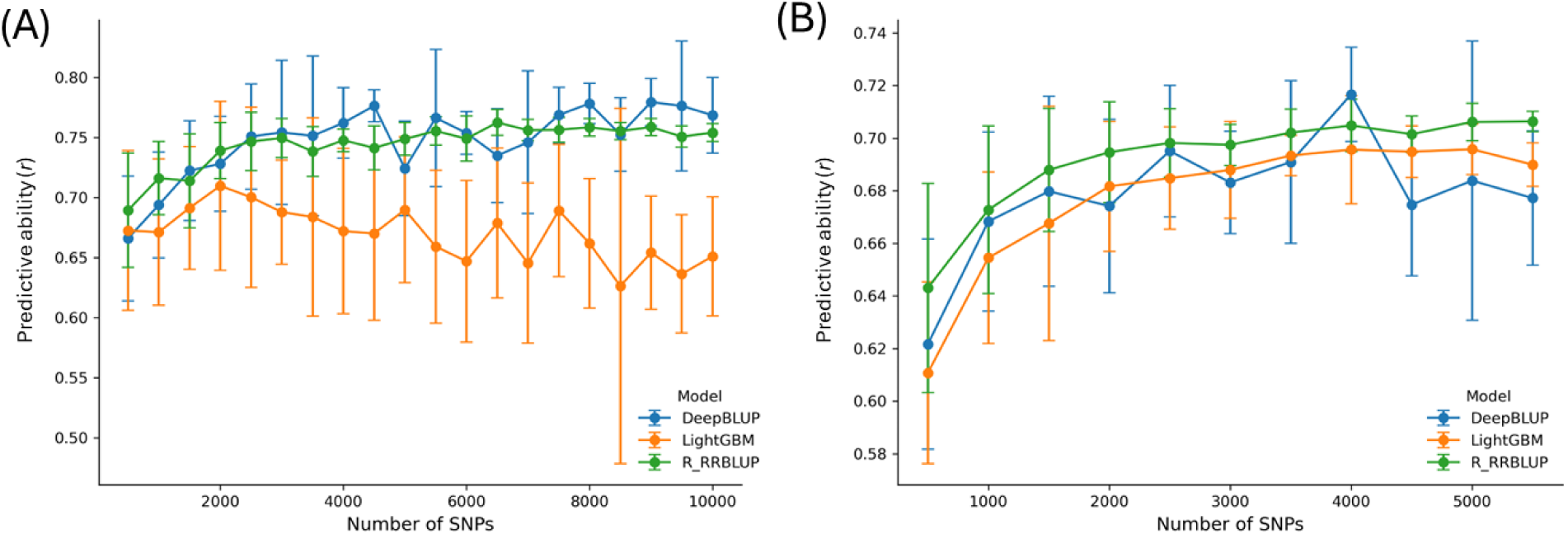
Predictive ability (PAs) of seed yield with increasing numbers of SNP markers. (A) BMEVSU260 test dataset, (B) BP295 test dataset. The CORE293 population was used as the training set, and predictions were generated using SNP-based R_RRBLUP (green), LightGBM (orange) and DeepBLUP (blue) models. SNP marker subsets of 500 to 10,000 (in 500 increments), were randomly sampled from the SNP set common to both training and test populations (x-axis). Ten independent random subsets were generated per SNP bins, and error bars represent standard deviation among replicates

Across both test populations, PA increased with marker density up to 2,000–2,500 SNPs, after which PA gains plateaued. However, at higher marker densities (>4,000 SNPs), we observed increased variability and occasional declines in PA, particularly for the LightGBM model in BMEVSU260 (Fig. 4A) and DeepBLUP in BP295 (Fig. 4B). This suggests that while more markers theoretically provide higher resolution, they may also introduce noise or lead to overfitting in certain model architectures under APP conditions. For the BMEVSU260 biparental population, R_RRBLUP and DeepBLUP reached a stable PA of approximately 0.74–0.76 at 2,500 SNPs. In contrast, LightGBM showed lower overall performance and a downward trend at high densities, likely due to the sensitivity of gradient boosting to high-dimensional feature spaces with limited training observations. In the BP295 population, while all three models peaked near ∼2,500 SNPs, DeepBLUP exhibited higher variance in the upper density range. Consequently, while 2,500 SNPs appear sufficient to capture the primary predictive signal, these results suggest that further increases in density do not merely yield diminishing returns but may introduce instability.

## Discussion

### Training population design for genomic prediction of seed yield

Training population composition emerged as a primary determinant of genomic prediction performance for seed yield in this study, particularly under APP scenarios that reflect actual breeding deployment. While historical core collections provide broad genetic diversity, our results demonstrate that genetic relevance to contemporary breeding germplasm outweighs population size or diversity alone for predicting the performance of new breeding lines.

The breeding-oriented population BP295 consistently outperformed historical CORE collections when predicting modern yellow- and brown-seeded breeding lines (YS38 and BS61), despite its lower marker density and narrower genetic diversity than CORE293 and CORE378. In contrast, CORE collections performed best when predicting genetically related biparental materials but showed reduced predictive ability when applied to elite breeding lines. These patterns indicate that broad germplasm diversity can dilute predictive signal when the training population is misaligned with breeding targets.

Notably, these contrasts were largely obscured under cross-validation but became evident under APP, highlighting the limitations of CV for evaluating deployment-ready GS strategies. Collectively, these results support the use of breeding-aligned, dynamically updated training populations as a foundation for effective genomic selection of complex traits such as seed yield.

### Genomic selection models and marker representations

Across all across-population prediction (APP) scenarios, classical linear mixed models remained among the most robust and consistently performing approaches for predicting seed yield in flax. Regularized linear models and Bayesian implementations showed stable performance across training–test combinations, highlighting their continued suitability for routine breeding applications. In contrast, more complex machine learning and deep learning models exhibited greater sensitivity to training population composition, with performance varying across test populations. Hybrid models that integrate RR-BLUP with deep architectures achieved competitive results in several scenarios, but did not consistently outperform linear baselines, indicating that increased model complexity alone does not guarantee improved robustness under APP.

From a practical breeding perspective, linear models such as RR-BLUP and BRR provided consistently robust performance across populations and marker representations while requiring minimal computational resources and parameter tuning. Although more complex machine learning and deep learning models occasionally achieved comparable or slightly improved predictive ability, their performance was less stable and more sensitive to training conditions. Therefore, linear models remain a reliable and efficient choice for routine GS implementation, while more complex models may be explored when sufficient computational resources and expertise are available.

Marker representations influenced prediction performance, although its effect was secondary to training population design. Across both cross-validation and APP analyses, SNP- and haplotype-based markers consistently outperformed PC markers, particularly when predicting contemporary breeding germplasm. PC markers frequently resulted in reduced or unstable prediction performance, especially for non-linear models, likely because dimensionality reduction obscures local LD patterns and additive genetic effects that are critical for yield prediction (Weber et al. 2023; Lin et al. 2024; Crossa et al. 2017; Du et al. 2018). By contrast, SNPs and haplotypes captured genome-wide LD more directly and provided broadly comparable performance across models. Haplotype-based markers achieved predictive abilities that were broadly comparable to those obtained with SNP markers, accounting for approximately half of the top-performing model–population combinations, while reducing marker dimensionality.

Together, these results indicate that robust genomic prediction of seed yield in flax depends more strongly on the genetic relevance of the training population and the use of genome-wide marker representations than on the choice of increasingly complex model architectures. Well-regularized linear models combined with moderate-density SNP or haplotype markers therefore remain reliable and practical choices for genomic selection in applied flax breeding programs.

### Genotyping-by-sequencing (GBS) markers

High-density SNP arrays are widely used in crops such as wheat, maize, and soybean (Lozada et al. 2019; Polzer et al. 2025; Li et al. 2024); however, GBS remains a primary genotyping platform for GS in many breeding programs due to its flexibility, scalability, and cost-effectiveness. In actual GS pipelines, training populations are often genotyped with high marker density (e.g., whole-genome sequencing or deep GBS), whereas test populations are genotyped at lower depth to reduce costs. This imbalance reduces the number of high-quality SNPs shared between training and test sets, which can directly influence APP performance.

Differences in SNP marker sets across populations, particularly between WGS- and GBS-derived datasets, may influence predictive ability in GS. Variability in marker density, genome coverage, and marker overlap between training and test populations can affect model performance. Although imputation and filtering were applied to harmonize datasets, incomplete marker overlap remains a potential limitation for across-population prediction. Standardized genotyping platforms or optimized marker panels could improve transferability across breeding populations.

Consistent with this expectation, APP predictive ability in our study was strongly associated with the number and genomic distribution of SNPs shared between training and test populations, and not strictly with the marker density *per se*. The combined biparental population BMEVSU260, which shared more than 35,000 SNPs with the training population, achieved PAs of up to r ∼ 0.83, comparable to within-population cross-validation results. In contrast, breeding populations genotyped using standard GBS protocols yielded approximately 12,000 SNPs, of which ∼7,000 were shared with the training set, resulting in predictive abilities up to r ∼ 0.79 across models and marker representations. These results indicate that once shared markers adequately capture genome-wide linkage disequilibrium, additional markers provide diminishing returns for PA.

This conclusion was further supported by marker subsampling simulations. Across two representative test populations (BMEVSU260 and BP295) and three model classes, PA increased rapidly with marker number up to 2,000–2,500 SNPs, beyond which gains plateaued (Fig. 4). Below this threshold, predictive abilities declined markedly, particularly for breeding populations with greater genetic heterogeneity. Together, these results suggest that moderate-density GBS panels (∼2,500–3,000 well-distributed SNPs) are sufficient to achieve stable and reliable PAs for seed yield in flax under APP conditions.

From a breeding perspective, these findings support the use of cost-efficient GBS strategies that prioritize an adequate number of shared markers over a larger number of markers. Such designs enable reliable genomic prediction while minimizing genotyping costs and computation time, thereby facilitating routine GS implementation for early-stage selection and optimizing resource allocation in flax breeding programs.

### Genomic selection integration into flax breeding program

APP remains one of the major challenges for implementing GS in practical breeding programs. While within-population CV often yields high predictive ability (You et al. 2022; Lan et al. 2020; He et al. 2019), its utility for predicting new breeding lines is limited. CV-based predictive abilities can be inflated due to shared genetic structure, close relatedness between training and test sets, and the ability of dense marker coverage to capture existing LD patterns. As a result, CV performance frequently overestimates the robustness of GS models which does not translate when applied to novel breeding germplasm.

Consistent with this limitation, relatively few studies have reported APP performance, and those that have generally observed lower PAs. For example, in wheat, PAs of grain yield ranged from -0.14 to 0.43 when two biparental populations were predicted using a diverse panel of soft red winter wheat accessions as the training set (Lozada et al. 2019). In CIMMYT’s elite yield nurseries, PAs declined from 0.35–0.44 within nurseries to 0.05–0.15 across nurseries, largely reflecting the genetic relatedness among lines (Juliana et al. 2018). Nevertheless, even under these conditions, GS at moderate selection intensity successfully identified a large proportion of top- performing lines (Juliana et al. 2018). Further improvements in APP have been achieved by expanding training populations across years and environments, with predictive ability increasing from 0.11 to 0.23 when five years of historical data were used for training (Vitale et al. 2025). Similar moderate APP predictive abilities have been reported for yield and quality traits in soybean and maize (Polzer et al. 2025; Dzievit et al. 2021).

Against this backdrop, the APP results obtained in this study demonstrate the operational potential of GS for flax breeding. When training and test populations were genetically aligned, GS achieved predictive abilities well within—or exceeding—the range considered useful for breeding decisions. For example, predicting the biparental population BMEVSU260 using CORE293 as the training population yielded predictive abilities as high as *r* = 0.83 with haplotype-based MLPGS (Fig. 3A). Similarly, using BP295 as the training population to predict YS38, haplotype-based GBLUP achieved *r* = 0.84 (Fig. 3B). For BS61, predictive ability reached *r* = 0.67 using HAP markers and LightGBM (Fig. 3C), while prediction of BP295 using CORE293, SNP markers and DNNGS achieved *r* = 0.73. These values exceed the commonly cited thresholds of practical utility for complex traits in cereals and legumes (*r* ∼ 0.3–0.6) (Crossa et al. 2017; Juliana et al. 2020). Notably, the relatively high APP predictive abilities observed here contrast with many previous reports for yield traits and appear to be driven primarily by training population alignment rather than by increased model complexity.

Predictive ability (PA) is not directly equivalent to the accuracy of breeding values because phenotypic observations include environmental variance. In contrast, phenotypic selection in breeding programs is typically based on multi-environment trials, which increases entry-mean heritability and improves selection reliability. Therefore, comparisons between GS and phenotypic selection should be interpreted in terms of decision consistency rather than direct equivalence of accuracy metrics. Thus, GS in this study was not evaluated solely as a prediction tool, but as a decision-support mechanism. GS-based selection consistently reproduced check-based advancement decisions while substantially reducing the number of lines requiring multi-location and multi-year field evaluation. By acting as an early-stage filter, GS enabled reductions of 60–90% in trial population size without increasing the risk of discarding elite candidates, thereby lowering costs and accelerating breeding cycles.

Based on these results, three practical principles emerge for integrating GS into flax breeding programs: (1) Training populations should be genetically and environmentally aligned with breeding targets, rather than relying exclusively on broad germplasm collections; (2) Training sets should be regularly updated with new breeding germplasm to maintain relevance and mitigate genetic drift; and (3) GS should be implemented as a gatekeeper in early selection stages, enabling substantial reductions in trial size (∼60–90%) once predictive abilities reach r ∼ 0.6–0.8, thereby focusing the program’s costly phenotyping resources on the most promising candidates.

Together, these findings indicate that GS for seed yield in flax is sufficiently robust for routine integration into breeding pipelines, particularly when used to complement—rather than replace—phenotypic evaluation.

### Cost of genomic selection

In flax breeding, field evaluation for seed yield represents a major financial and logistical investment. In the CDC flax breeding program, the cost per field plot—from seeding through harvest and data collection—is approximately $75 CAD, resulting in $150 per line for two replications at a single location in preliminary yield trials. Additional routine evaluations further increase costs, including oil profile determination by gas chromatography (FAME; ∼$10 per sample) and disease resistance screening (e.g., Fusarium wilt; ∼$12.50 per row). With two replications per line, the total cost per line is therefore approximately $195 CAD, and advancing 300 lines to preliminary trials requires an annual investment of approximately $58,500 CAD.

In contrast, the per-sample cost of genotyping-by-sequencing (GBS) is substantially lower. DNA preparation and library construction together cost approximately $17.10 per sample, while sequencing costs depend on sequencing platform and sample throughput. On the NovaSeq platform, sequencing costs scaled linearly with read depth and sample multiplexing. Across scenarios ranging from 192 to 768 samples per run, the per-sample sequencing costs remain linear. For example, using 768 samples on the NovaSeq 3200M PE150 platform, the per-sample cost can be approximated as *y = 1.3594x + 17.121*, where *y* is the cost per sample (CAD) and *x* is the number of million PE150 reads targeted. Under this model, a sequencing depth of 6M PE150 reads per sample—corresponding to approximately 6,000 common SNPs—results in a per-sample cost of ∼$25.27 CAD. Based on the results from the marker number simulations (Fig. 4), ∼2,500 genome-wide SNPs are likely sufficient to achieve stable and near-maximal PA for seed yield across breeding populations, the ∼6,000 SNPs generated at this sequencing depth therefore exceed the minimum marker requirement, providing a conservative and robust genotyping strategy. Under this scenario, the total GBS cost for 300 lines is approximately $7,581 CAD. Based on check-based selection analyses, reducing the number of lines advanced to preliminary and subsequent yield trials by 61–91% would decrease the initial pool of 300 lines to 27–117 lines. Under this strategy, the combined cost of genotyping, yield trials, disease screening, and FAME analysis is estimated at $12,846–$30,396 CAD, corresponding to 48–78% cost savings relative to phenotypic selection alone.

Overall, these results demonstrate that GS can reduce the cost and scale of field evaluations conducted at early breeding stages (e.g., preliminary yield trials following initial line development), while retaining high-performing candidate lines. In addition to direct cost savings, earlier GS-based selection enables advancement of selected lines to contra-season nurseries (e.g., winter nurseries in New Zealand), further reducing breeding cycle time. By enabling earlier selection decisions, smaller field trials, and more efficient resource allocation, GS offers a practical and cost-effective strategy for accelerating flax breeding without increasing the risk of discarding elite germplasm.

While the cost analysis presented here reflects the initial integration of GS, maintaining PA over successive breeding cycles requires periodic updating of the training population with newly phenotyped breeding lines to mitigate model decay. This model upkeep incurs recurrent costs associated with genotyping and limited phenotyping; however, these activities are typically integrated into routine breeding operations and represent a small fraction of the extensive multi-location field-testing costs that are reduced by GS. Although ongoing investment in training population renewal is necessary, the net economic advantage of GS-assisted breeding remains strongly positive.

## Conclusion

This study demonstrates that genomic selection for seed yield in flax can be effectively deployed under realistic breeding scenarios when training populations are genetically aligned with breeding targets. Across independent test populations, training population composition emerged as the primary determinant of predictive ability, often outweighing differences among models or marker representations. Genomic selection consistently reproduced check-based advancement decisions while substantially reducing the number of lines requiring extensive field evaluation, indicating its value as an early-stage decision-support tool rather than a replacement for phenotypic selection. Moderate-density GBS marker panels were sufficient to achieve stable predictive ability, enabling cost-efficient implementation. Together, these results provide practical guidance for integrating genomic selection into flax breeding programs and highlight the importance of breeding-oriented training population design for achieving robust prediction of complex traits. When applied as a complement to phenotypic evaluation, genomic selection offers a viable pathway to improve breeding efficiency, reduce costs, and accelerate cultivar development.

## Supplementary Information

The online version contains supplementary material available at xxxx.

## Acknowledgements

We thank Dr. Liqian He for reviewing the draft manuscript and for providing valuable comments and suggestions, and Tara Edwards for DNA preparation.

## Author contribution statement

FY contributed to conceptualization, funding acquisition, investigation, methodology, data analysis, and original draft preparation. SC contributed to conceptualization, funding acquisition, DNA preparation for SNP genotyping, data curation, and manuscript review and editing. BT contributed to conceptualization, funding acquisition, data analysis, and manuscript review and editing. JD contributed to software development. CZ, JD, and PL contributed to formal analysis, visualization, data curation, and manuscript review. KJ, MH and BT performed field trials for breeding lines and manuscript review. All authors read and approved the final manuscript.

## Funding

This Sustainable Canadian Agricultural Partnership AgriScience Cluster (SCAP-ASC) project Activity 5A (SCAP-ACS-05) was funded by Agriculture and Agri-Food Canada (AAFC) and the Diverse Field Crop Cluster (DFCC) managed by Ag-West Bio Inc. The core collection research originated from the Total Utilization Flax GENomics (TUFGEN) project funded by Genome Canada and affiliated stakeholders.

## Declarations

## Conflict of interest

The authors declare they have no conflict of interest.

## Supplementary Tables and Figures

**Fig. S1.**
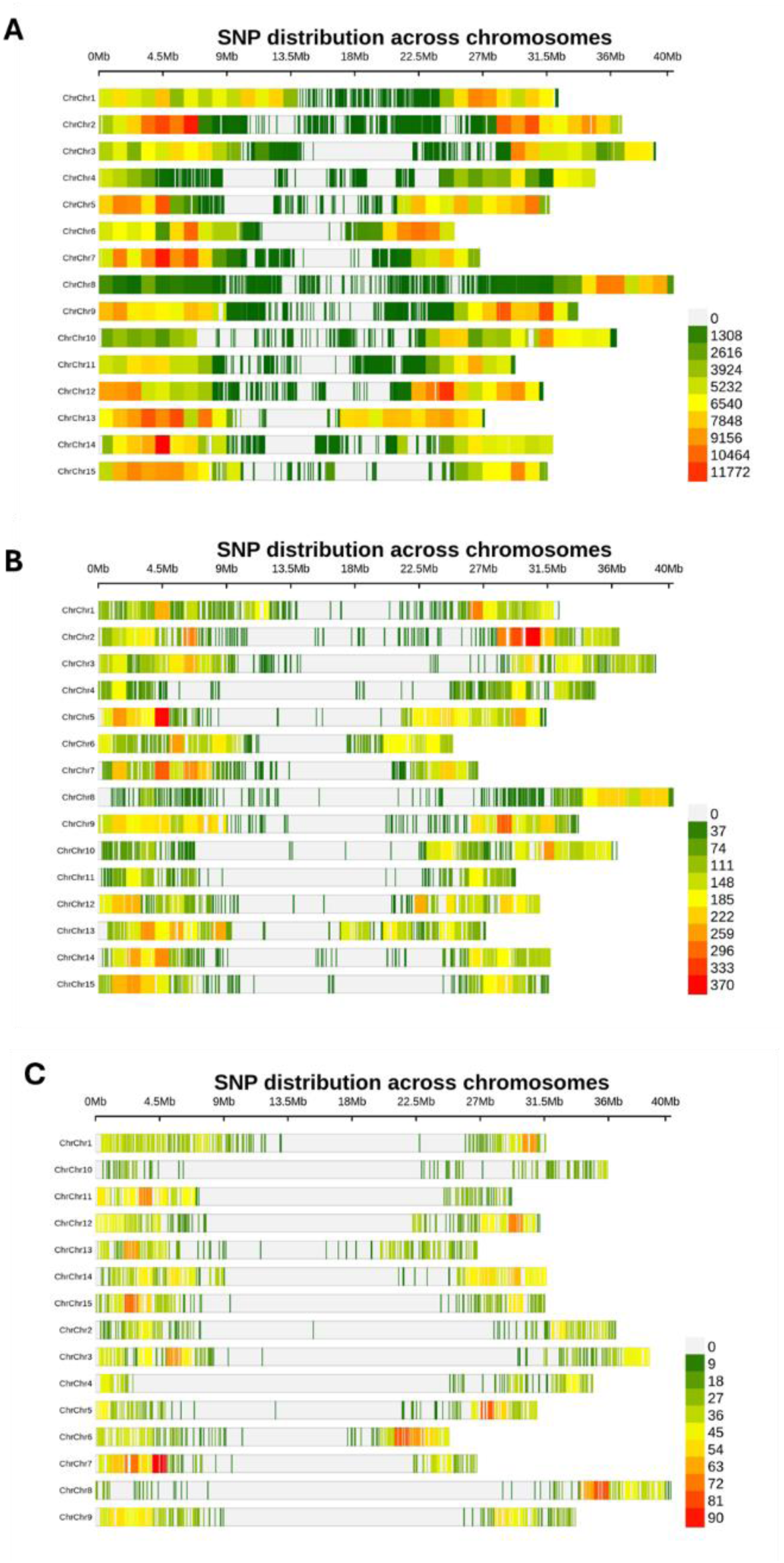
SNP density along chromosomes (A) in the core collection of 378 accessions (CORE378) comprising ∼1.7M SNPs, (B) 33,596 SNPs shared between CORE378 and the 260 test lines from the combined biparental population BMEVSU260, and (C) 5,604 SNPs shared by CORE378, CORE293, YS38, BS61 and BP295. Plots were generated with a 100-Kb window using the CMplot R package (Yin et al. 2021)

**Fig. S2.**
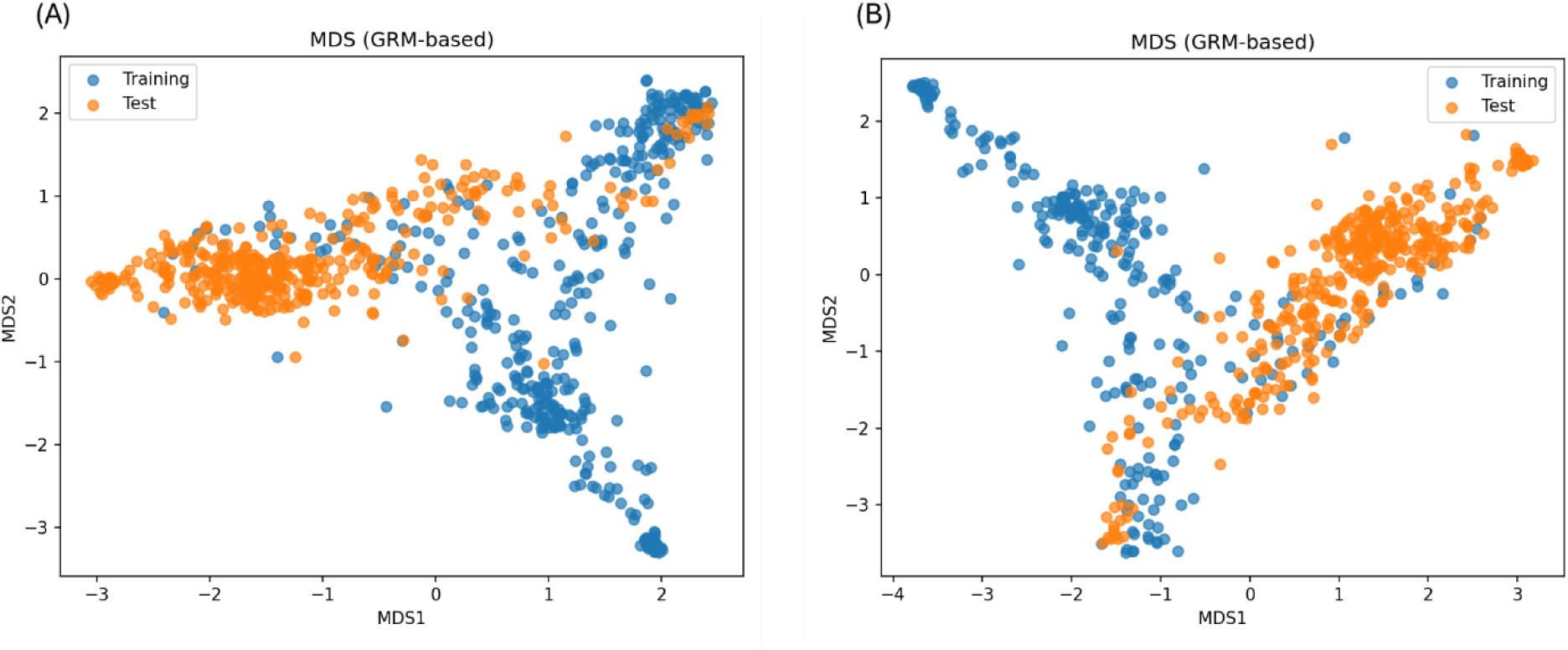
Genetic relationships between training and test populations based on GRM-derived multidimensional scaling (MDS). (A) CORE378 (training; blue) versus BP295 (test; orange). (B) CORE293 (training; blue) versus BP295 (test; orange). Dots represent individual genotypes projected onto the first two MDS axes derived from the genomic relationship matrix (GRM). In panel (A), CORE378 shows broader genetic coverage and greater overlap with BP295, whereas in panel (B), CORE293 exhibits more limited overlap with BP295

**Table S1.**
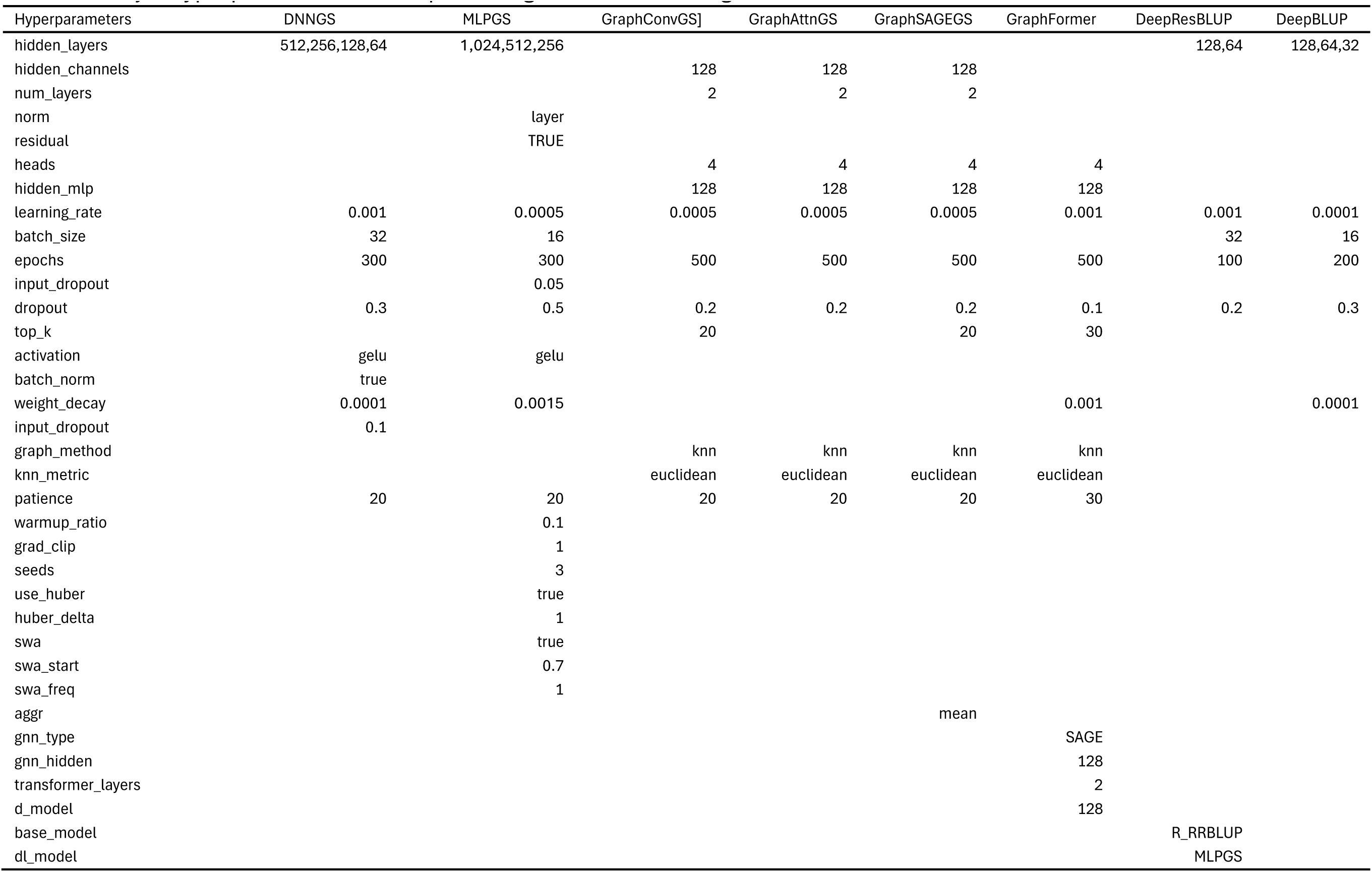
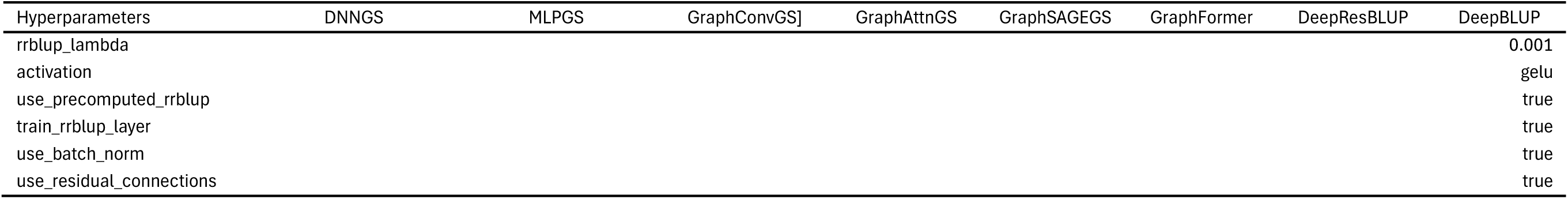
Major hyperparameters of deep learning models used for genomic selection.

**Table S2.**
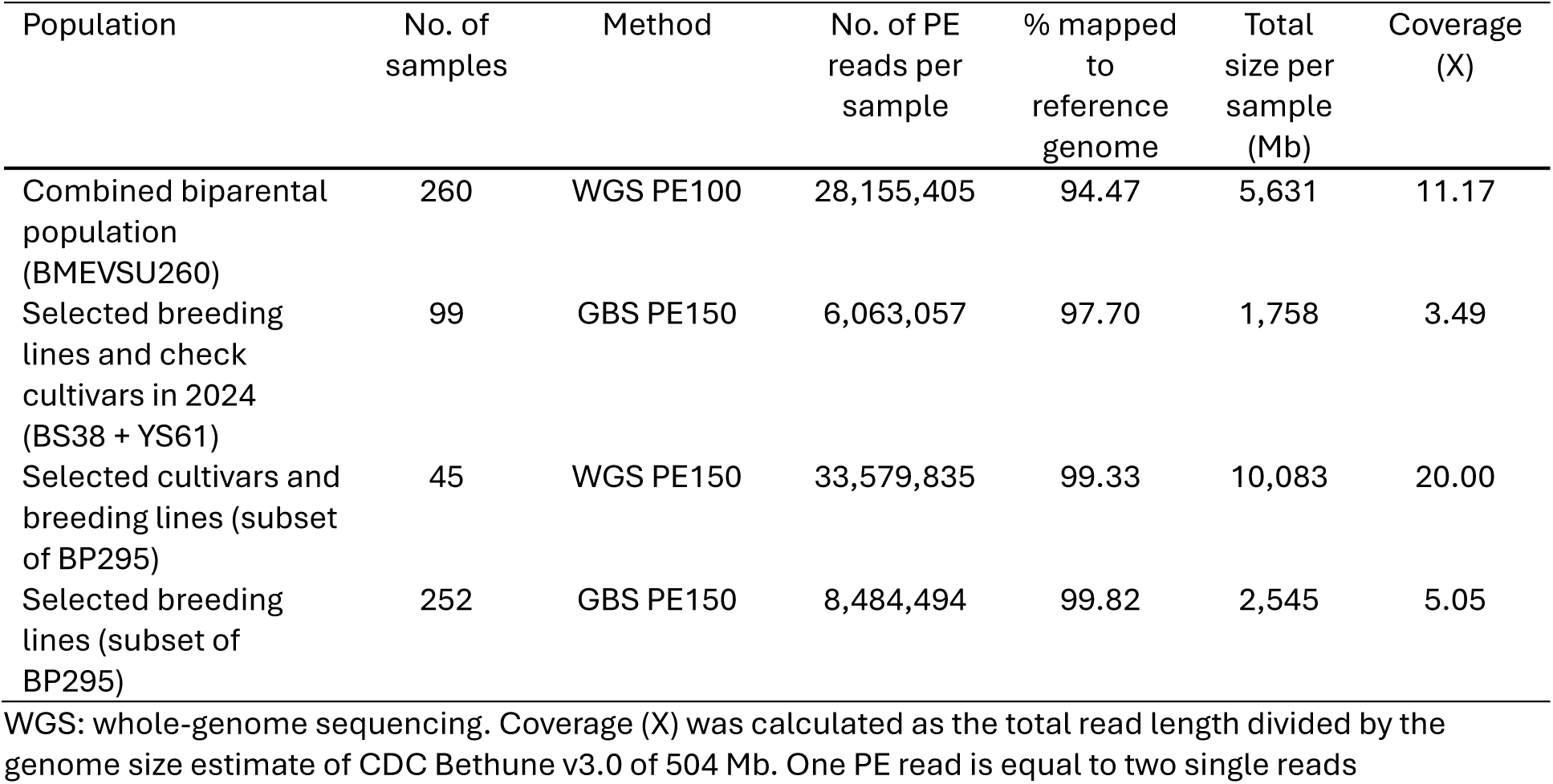
Summary statistics of Illumina paired-end (PE) reads generated for the test populations.

**Table S3.** Predictive abilities (PAs) and their standard deviations for seed yield (YLD) under five-fold cross-validation (CV) using BP295, CORE293 and CORE378.

| Model | BP295 |  |  |  | CORE293 |  |  |  | CORE378 |  |  |  |
| --- | --- | --- | --- | --- | --- | --- | --- | --- | --- | --- | --- | --- |
|  | HAP | PC | SNP | Max | HAP | PC | SNP | Max | HAP | PC | SNP | Max |
| R_RRLUP | 0.840 ± | 0.750 ± | 0.837 ± | 0.840 ± | 0.840 ± | 0.834 ± | 0.832 ± | 0.840 ± | 0.819 ± | 0.815 ± | 0.814 ± | 0.819 ± |
|  | 0.038 | 0.145 | 0.038 | 0.038 | 0.029 | 0.030 | 0.030 | 0.029 | 0.042 | 0.042 | 0.043 | 0.042 |
| R_GBLUP | 0.787 ± | 0.463 ± | 0.781 ± | 0.787 ± | 0.738 ± | 0.528 ± | 0.736 ± | 0.738 ± | 0.663 ± | 0.624 ± | 0.662 ± | 0.663 ± |
|  | 0.052 | 0.169 | 0.054 | 0.052 | 0.052 | 0.090 | 0.052 | 0.052 | 0.070 | 0.110 | 0.070 | 0.070 |
| BRR | 0.839 ± | 0.721 ± | 0.812 ± | 0.839 ± | 0.840 ± | 0.834 ± | 0.832 ± | 0.840 ± | 0.818 ± | 0.814 ± | 0.814 ± | 0.818 ± |
|  | 0.039 | 0.149 | 0.056 | 0.039 | 0.029 | 0.030 | 0.030 | 0.029 | 0.042 | 0.042 | 0.043 | 0.042 |
| ElasticNet | 0.814 ± | 0.744 ± | 0.811 ± | 0.814 ± | 0.803 ± | 0.844 ± | 0.806 ± | 0.844 ± | 0.789 ± | 0.819 ± | 0.797 ± | 0.819 ± |
|  | 0.042 | 0.146 | 0.047 | 0.042 | 0.034 | 0.028 | 0.035 | 0.028 | 0.047 | 0.041 | 0.045 | 0.041 |
| RFR | 0.799 ± | 0.681 ± | 0.803 ± | 0.803 ± | 0.812 ± | 0.797 ± | 0.815 ± | 0.815 ± | 0.791 ± | 0.776 ± | 0.799 ± | 0.799 ± |
|  | 0.052 | 0.246 | 0.048 | 0.048 | 0.036 | 0.039 | 0.036 | 0.036 | 0.043 | 0.044 | 0.040 | 0.040 |
| LightGBM | 0.793 ± | 0.482 ± | 0.789 ± | 0.793 ± | 0.810 ± | 0.772 ± | 0.822 ± | 0.822 ± | 0.779 ± | 0.751 ± | 0.792 ± | 0.792 ± |
|  | 0.054 | 0.161 | 0.045 | 0.054 | 0.040 | 0.052 | 0.038 | 0.038 | 0.045 | 0.049 | 0.047 | 0.047 |
| XGBoost | 0.779 ± | 0.615 ± | 0.758 ± | 0.779 ± | 0.781 ± | 0.775 ± | 0.788 ± | 0.788 ± | 0.753 ± | 0.759 ± | 0.764 ± | 0.764 ± |
|  | 0.051 | 0.243 | 0.063 | 0.051 | 0.049 | 0.053 | 0.047 | 0.047 | 0.048 | 0.043 | 0.048 | 0.048 |
| MLPGS | 0.790 ± | 0.693 ± | 0.794 ± | 0.794 ± | 0.825 ± | 0.804 ± | 0.811 ± | 0.825 ± | 0.803 ± | 0.798 ± | 0.802 ± | 0.803 ± |
|  | 0.048 | 0.181 | 0.053 | 0.053 | 0.037 | 0.043 | 0.050 | 0.037 | 0.050 | 0.044 | 0.045 | 0.050 |
| DNNGS | 0.808 ± | 0.507 ± | 0.823 ± | 0.823 ± | 0.815 ± | 0.806 ± | 0.799 ± | 0.815 ± | 0.810 ± | 0.776 ± | 0.804 ± | 0.810 ± |
|  | 0.063 | 0.347 | 0.050 | 0.050 | 0.051 | 0.041 | 0.074 | 0.051 | 0.046 | 0.083 | 0.052 | 0.046 |
| GraphConvGS | 0.703 ± | 0.252 ± | 0.638 ± | 0.703 ± | 0.613 ± | 0.594 ± | 0.594 ± | 0.613 ± | 0.557 ± | 0.521 ± | 0.535 ± | 0.557 ± |
|  | 0.122 | 0.267 | 0.189 | 0.122 | 0.055 | 0.144 | 0.092 | 0.055 | 0.082 | 0.129 | 0.102 | 0.082 |
| GraphAttnGS | 0.699 ± | 0.476 ± | 0.631 ± | 0.699 ± | 0.547 ± | 0.463 ± | 0.551 ± | 0.551 ± | 0.531 ± | 0.447 ± | 0.471 ± | 0.531 ± |
|  | 0.081 | 0.244 | 0.170 | 0.081 | 0.153 | 0.255 | 0.145 | 0.145 | 0.105 | 0.183 | 0.226 | 0.105 |
| GraphSAGEGS | 0.765 ± | 0.724 ± | 0.760 ± | 0.765 ± | 0.807 ± | 0.750 ± | 0.783 ± | 0.807 ± | 0.772 ± | 0.731 ± | 0.763 ± | 0.772 ± |
|  | 0.065 | 0.146 | 0.060 | 0.065 | 0.039 | 0.055 | 0.042 | 0.039 | 0.047 | 0.055 | 0.053 | 0.047 |
| GraphFormer | 0.734 ± | 0.728 ± | 0.730 ± | 0.734 ± | 0.817 ± | 0.783 ± | 0.796 ± | 0.817 ± | 0.780 ± | 0.758 ± | 0.770 ± | 0.780 ± |
|  | 0.077 | 0.151 | 0.071 | 0.077 | 0.035 | 0.039 | 0.039 | 0.035 | 0.048 | 0.045 | 0.047 | 0.048 |
| DeepResBLUP | 0.829 ± | 0.737 ± | 0.828 ± | 0.829 ± | 0.836 ± | 0.813 ± | 0.829 ± | 0.836 ± | 0.813 ± | 0.791 ± | 0.812 ± | 0.813 ± |
|  | 0.042 | 0.148 | 0.040 | 0.042 | 0.032 | 0.036 | 0.034 | 0.032 | 0.043 | 0.043 | 0.044 | 0.043 |
| DeepBLUP | 0.821 ± | 0.726 ± | 0.805 ± | 0.821 ± | 0.828 ± | 0.826 ± | 0.815 ± | 0.828 ± | 0.805 ± | 0.803 ± | 0.797 ± | 0.805 ± |
|  | 0.050 | 0.156 | 0.054 | 0.050 | 0.030 | 0.031 | 0.038 | 0.030 | 0.044 | 0.043 | 0.047 | 0.044 |
| EnsembleGS | 0.794 ± | 0.709 ± | 0.792 ± | 0.794 ± | 0.807 ± | 0.771 ± | 0.819 ± | 0.819 ± | 0.775 ± | 0.747 ± | 0.785 ± | 0.785 ± |
|  | 0.055 | 0.156 | 0.047 | 0.055 | 0.039 | 0.051 | 0.038 | 0.038 | 0.042 | 0.048 | 0.047 | 0.047 |

**Table S4.** Predictive abilities (PAs) of four test populations for seed yield (YLD) under across-population prediction (APP) using CORE293 as the training population.

| Model | Marker representation | Test population |  |  |  |
| --- | --- | --- | --- | --- | --- |
|  |  | BP295 | YS38 | BS61 | BMEVSU260 |
| R_RRBLUP | PC | 0.707 | 0.647 | 0.543 | 0.772 |
|  | HAP | 0.717 | 0.585 | 0.595 | 0.794 |
|  | SNP | 0.707 | 0.624 | 0.558 | 0.762 |
| R_GBLUP | PC | 0.586 | 0.711 | 0.521 | 0.310 |
|  | HAP | 0.522 | 0.681 | 0.592 | 0.694 |
|  | SNP | 0.499 | 0.711 | 0.521 | 0.314 |
| BRR | PC | 0.707 | 0.647 | 0.542 | 0.772 |
|  | HAP | 0.717 | 0.584 | 0.595 | 0.794 |
|  | SNP | 0.707 | 0.624 | 0.558 | 0.761 |
| ElasticNet | PC | 0.710 | 0.668 | 0.559 | 0.774 |
|  | HAP | 0.690 | 0.571 | 0.616 | 0.722 |
|  | SNP | 0.699 | 0.526 | 0.578 | 0.723 |
| RFR | PC | 0.680 | 0.658 | 0.370 | 0.738 |
|  | HAP | 0.717 | 0.470 | 0.574 | 0.681 |
|  | SNP | 0.716 | 0.393 | 0.363 | 0.627 |
| LightGBM | PC | 0.731 | 0.586 | 0.299 | 0.763 |
|  | HAP | 0.673 | 0.528 | 0.598 | 0.621 |
|  | SNP | 0.674 | 0.400 | 0.434 | 0.626 |
| XGBoost | PC | 0.696 | 0.542 | 0.249 | 0.679 |
|  | HAP | 0.692 | 0.536 | 0.466 | 0.647 |
|  | SNP | 0.612 | 0.451 | 0.440 | 0.595 |
| MLPGS | PC | 0.691 | 0.352 | 0.251 | 0.781 |
|  | HAP | 0.730 | 0.267 | 0.095 | 0.824 |
|  | SNP | 0.721 | 0.305 | 0.183 | 0.806 |
| DNNGS | PC | 0.693 | 0.698 | 0.517 | 0.777 |
|  | HAP | 0.709 | 0.590 | 0.546 | 0.808 |
|  | SNP | 0.734 | 0.648 | 0.546 | 0.789 |
| GraphConvGS | PC | 0.496 | 0.656 | -0.372 | 0.820 |
|  | HAP | 0.648 | 0.538 | -0.151 | 0.835 |
|  | SNP | 0.624 | 0.589 | -0.359 | 0.808 |
| GraphAttnGS | PC | 0.508 | 0.710 | -0.423 | 0.685 |
|  | HAP | 0.572 | 0.411 | -0.108 | 0.788 |
|  | SNP | 0.568 | 0.511 | 0.518 | 0.834 |
| GraphSAGEGS | PC | 0.646 | 0.303 | -0.005 | 0.779 |
|  | HAP | 0.719 | 0.128 | 0.284 | 0.829 |
|  | SNP | 0.728 | 0.223 | 0.188 | 0.800 |
| GraphFormer | PC | 0.697 | 0.306 | 0.168 | 0.754 |
|  | HAP | 0.718 | 0.034 | 0.171 | 0.822 |
|  | SNP | 0.692 | 0.259 | 0.196 | 0.817 |
|  |  | BP295 | YS38 | BS61 | BMEVSU260 |
| DeepResBLUP | PC | 0.655 | 0.574 | 0.347 | 0.781 |
|  | HAP | 0.696 | 0.598 | 0.598 | 0.789 |
|  | SNP | 0.710 | 0.648 | 0.535 | 0.784 |
| DeepBLUP | PC | 0.673 | 0.650 | 0.585 | 0.752 |
|  | HAP | 0.731 | 0.552 | 0.484 | 0.798 |
|  | SNP | 0.663 | 0.684 | 0.567 | 0.781 |
| EnsembleGS | PC | 0.728 | 0.546 | 0.236 | 0.776 |
|  | HAP | 0.702 | 0.450 | 0.553 | 0.742 |
|  | SNP | 0.653 | 0.307 | 0.380 | 0.690 |
| PC mean |  | 0.663 | 0.578 | 0.274 | 0.732 |
| HAP mean |  | 0.685 | 0.470 | 0.407 | 0.762 |
| SNP mean |  | 0.669 | 0.494 | 0.388 | 0.720 |
| PC max |  | 0.731 | 0.711 | 0.585 | 0.820 |
| HAP max |  | 0.731 | 0.681 | 0.616 | 0.835 |
| SNP max |  | 0.734 | 0.711 | 0.578 | 0.834 |

**Table S5.** Predictive abilities (PAs) of four test populations for seed yield (YLD) under across-population prediction (APP) using CORE378 as the training population.

| Model | Marker representation | Test population |  |  |  |
| --- | --- | --- | --- | --- | --- |
|  |  | BP295 | YS38 | BS61 | BMEVSU260 |
| R_RRLUP | PC | 0.720 | 0.598 | 0.451 | 0.703 |
|  | HAP | 0.737 | 0.446 | 0.500 | 0.704 |
|  | SNP | 0.728 | 0.600 | 0.493 | 0.697 |
| R_GBLUP | PC | 0.664 | 0.664 | 0.494 | 0.789 |
|  | HAP | 0.673 | 0.608 | 0.565 | 0.765 |
|  | SNP | 0.664 | 0.664 | 0.494 | 0.790 |
| BRR | PC | 0.720 | 0.597 | 0.451 | 0.703 |
|  | HAP | 0.737 | 0.446 | 0.500 | 0.704 |
|  | SNP | 0.728 | 0.599 | 0.493 | 0.697 |
| ElasticNet | PC | 0.726 | 0.626 | 0.444 | 0.749 |
|  | HAP | 0.737 | 0.485 | 0.588 | 0.708 |
|  | SNP | 0.732 | 0.490 | 0.555 | 0.701 |
| RFR | PC | 0.740 | 0.554 | 0.435 | 0.612 |
|  | HAP | 0.716 | 0.247 | 0.553 | 0.673 |
|  | SNP | 0.728 | 0.224 | 0.531 | 0.694 |
| LightGBM | PC | 0.717 | 0.595 | 0.500 | 0.652 |
|  | HAP | 0.716 | 0.231 | 0.535 | 0.673 |
|  | SNP | 0.696 | 0.245 | 0.579 | 0.691 |
| XGBoost | PC | 0.700 | 0.202 | 0.438 | 0.686 |
|  | HAP | 0.675 | 0.214 | 0.553 | 0.586 |
|  | SNP | 0.668 | 0.435 | 0.516 | 0.672 |
| MLPGS | PC | 0.735 | 0.320 | 0.267 | 0.645 |
|  | HAP | 0.732 | 0.271 | 0.181 | 0.596 |
|  | SNP | 0.744 | 0.380 | 0.009 | 0.609 |
| DNNGS | PC | 0.707 | 0.623 | 0.572 | 0.715 |
|  | HAP | 0.767 | 0.515 | 0.481 | 0.758 |
|  | SNP | 0.752 | 0.504 | 0.544 | 0.680 |
| GraphConvGS | PC | 0.533 | 0.289 | -0.655 | 0.658 |
|  | HAP | 0.422 | 0.528 | -0.350 | 0.824 |
|  | SNP | 0.473 | 0.531 | -0.583 | 0.842 |
| GraphAttnGS | PC | 0.587 | 0.245 | -0.268 | 0.662 |
|  | HAP | 0.561 | 0.005 | -0.276 | 0.833 |
|  | SNP | 0.596 | -0.271 | 0.233 | 0.772 |
| GraphSAGEGS | PC | 0.685 | 0.338 | 0.243 | 0.698 |
|  | HAP | 0.742 | 0.091 | 0.145 | 0.743 |
|  | SNP | 0.742 | 0.181 | 0.140 | 0.747 |
| GraphFormer | PC | 0.687 | 0.325 | 0.304 | 0.406 |
|  | HAP | 0.735 | 0.118 | 0.047 | 0.735 |
|  | SNP | 0.726 | 0.161 | 0.018 | 0.719 |
|  |  | BP295 | YS38 | BS61 | BMEVSU260 |
| DeepResBLUP | PC | 0.731 | 0.565 | 0.525 | 0.653 |
|  | HAP | 0.713 | 0.446 | 0.375 | 0.713 |
|  | SNP | 0.745 | 0.580 | 0.470 | 0.639 |
| DeepBLUP | PC | 0.665 | 0.552 | 0.436 | 0.696 |
|  | HAP | 0.737 | 0.407 | 0.514 | 0.687 |
|  | SNP | 0.763 | 0.570 | 0.528 | 0.673 |
| EnsembleGS | PC | 0.711 | 0.575 | 0.496 | 0.637 |
|  | HAP | 0.706 | 0.190 | 0.527 | 0.650 |
|  | SNP | 0.675 | 0.187 | 0.570 | 0.684 |
| PC mean |  | 0.689 | 0.479 | 0.321 | 0.667 |
| HAP mean |  | 0.694 | 0.328 | 0.340 | 0.710 |
| SNP mean |  | 0.698 | 0.380 | 0.349 | 0.707 |
| PC max |  | 0.740 | 0.664 | 0.572 | 0.789 |
| HAP max |  | 0.767 | 0.608 | 0.588 | 0.833 |
| SNP max |  | 0.763 | 0.664 | 0.579 | 0.842 |

**Table S6.** Predictive abilities (PAs) of two test populations for seed yield (YLD) under across-population prediction (APP) using BP295 as the training population.

| Model | Marker representation | YS38 | BS61 |
| --- | --- | --- | --- |
| R_RRLUP | PC | 0.726 | 0.570 |
|  | HAP | 0.731 | 0.508 |
|  | SNP | 0.728 | 0.571 |
| R_GBLUP | PC | 0.439 | 0.503 |
|  | HAP | 0.841 | 0.652 |
|  | SNP | 0.848 | 0.665 |
| BRR | PC | 0.725 | 0.568 |
|  | HAP | 0.730 | 0.504 |
|  | SNP | 0.453 | 0.475 |
| ElasticNet | PC | 0.717 | 0.610 |
|  | HAP | 0.674 | 0.666 |
|  | SNP | 0.654 | 0.659 |
| RFR | PC | 0.424 | 0.125 |
|  | HAP | 0.555 | 0.564 |
|  | SNP | 0.635 | 0.664 |
| LightGBM | PC | 0.595 | 0.076 |
|  | HAP | 0.755 | 0.561 |
|  | SNP | 0.623 | 0.670 |
| XGBoost | PC | 0.513 | 0.290 |
|  | HAP | 0.383 | 0.450 |
|  | SNP | 0.462 | 0.457 |
| MLPGS | PC | 0.745 | 0.643 |
|  | HAP | 0.711 | 0.153 |
|  | SNP | 0.775 | 0.262 |
| DNNGS | PC | 0.740 | 0.588 |
|  | HAP | 0.273 | 0.668 |
|  | SNP | -0.076 | 0.136 |
| GraphConvGS | PC | 0.669 | 0.379 |
|  | HAP | 0.688 | -0.373 |
|  | SNP | 0.670 | 0.477 |
| GraphAttnGS | PC | -0.237 | 0.511 |
|  | HAP | -0.560 | 0.546 |
|  | SNP | 0.217 | 0.362 |
| GraphSAGEGS | PC | 0.720 | 0.589 |
|  | HAP | 0.726 | 0.272 |
|  | SNP | 0.769 | 0.214 |
| GraphFormer | PC | 0.729 | 0.319 |
|  | HAP | 0.774 | 0.380 |
|  | SNP | 0.701 | 0.242 |

| Model | Marker<br>representation | YS38 | BS61 |
| --- | --- | --- | --- |
| DeepResBLUP | PC | 0.798 | 0.539 |
|  | HAP | 0.614 | 0.386 |
|  | SNP | 0.633 | 0.474 |
| DeepBLUP | PC | 0.719 | 0.574 |
|  | HAP | 0.674 | 0.340 |
|  | SNP | 0.727 | 0.237 |
| EnsembleGS | PC | 0.720 | 0.453 |
|  | HAP | 0.790 | 0.500 |
|  | SNP | 0.525 | 0.530 |
| PC mean |  | 0.609 | 0.459 |
| HAP mean |  | 0.585 | 0.424 |
| SNP mean |  | 0.584 | 0.443 |
| PC max |  | 0.798 | 0.643 |
| HAP max |  | 0.841 | 0.668 |
| SNP max |  | 0.848 | 0.670 |

## Notes

### Competing Interest Statement

The authors have declared no competing interest.

### Summary of Updates

Training and test population names were changed.

